# A Bayesian framework for comparing the structure of spontaneous correlated activity recorded under different conditions

**DOI:** 10.1101/037358

**Authors:** Catherine S Cutts, Stephen J Eglen

## Abstract

Distant-dependent correlations in spontaneous retinal activity are thought to be instructive in the development of the retinotopic map and eye-specific segregation maps. Many studies which seek to investigate these correlations and their role in map formation record spontaneous retinal activity from different phenotypes or experimental conditions and compare the distance-dependence of the correlations between different conditions. They seek to demonstrate that these correlations differ significantly, and this analysis is often key to the study’s conclusions. In this work, we assess the methods of inference which have been previously used to investigate this problem and conclude that they are inadequate. We propose a hierarchical Bayesian framework to model distant-dependent correlations in spontaneous retinal activity data and specify a method which uses the data to specify the form of the model. This model allows us to assess the evidence for/against differences in correlations between experimental conditions in a more robust and credible way. We demonstrate the use of this method by applying it to data from two studies of spontaneous retinal activity. We believe however the framework to be rather more general and that it can be used in a wide range of datasets where distance and correlation are substitute for other independent and dependent variables from experiments.

## Introduction

During early development of the visual system, the retina generates spontaneous patterns of neuronal activity [22,40]. Pairs of retinal ganglion cells (RGCs) that are close to each other tend to have highly correlated activity, whereas activity in pairs of neurons that are a long distance apart (typically over 300-500 |j.m) is less correlated [33,66]. This distance-dependence in correlation between pairs of RGCs is thought to be a cue to help establish topographic maps [65] as when the distance-dependence correlations are perturbed (e.g. in a mutant mouse) topographic maps are perturbed [34].

The standard approach [66] used by investigators to assess distance-dependent correlations is to record spontaneous activity using a multielectrode array (MEA). For each pair of electrodes in a recording, we then plot the correlation in activity of the two spike trains as a function of distance separating the two electrodes. For an MEA with *N* spike trains, this generates *N*(*N* — 1)/2 datapoints which can be plotted individually (Figure 1A) or are commonly summarised by fitting an exponential decaying function to them (Figure 1B; [66]). By assessing spontaneous activity under two different experimental conditions (e.g. wild type vs mutant) we can then compare their resulting distance-dependence profiles. In the first example dataset shown in Figure 1B, the two curves seem quite different, leading us to conclude that the correlations are quite different. However, in Figure 1C, it is harder to discern whether the curves are distinct. Furthermore, sometimes, we may wish to compare more than two curves simultaneously (Figure 2D) when we have either more than two experimental conditions, or when there are multiple recordings from each condition.

**Figure 1:**
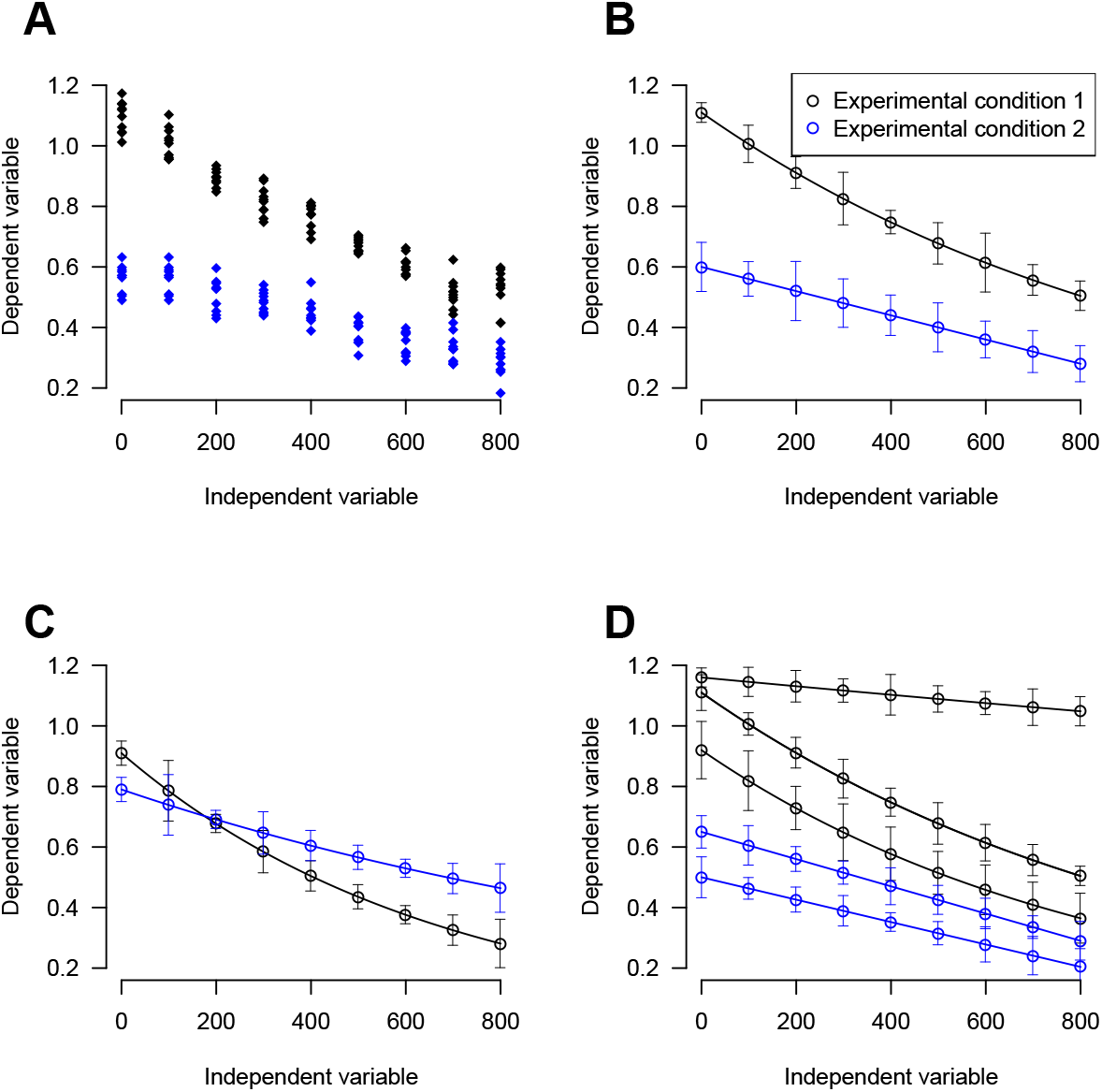
Determining if two experimental conditions differ based on inspection of graphical summaries is difficult and insufficiently rigorous. A: Synthetic data from two different experimental conditions. Ten measurements of a dependent variable are shown at nine different values of an independent variable for two experimental conditions. Is there a difference in the dependent variable between the two experimental conditions? B: Data from A summarised by mean (at each value of the independent variable) and ± one standard deviation (s.d.). Whilst this may aid visual clarity, information is lost and conclusions drawn on the basis of visual inspection alone may be misleading. Visual inspection of A and B shows that a difference in observations between the two conditions is likely since there is no overlap of points at any distance (and no overlap of error bars). It is difficult to judge whether there is a difference in the dependence on the independent variable between the two conditions. C: A different synthetic data set is summarised as per B, such that this time it is less obvious whether the two populations differ e.g. there is partial overlap of error bars. D: A third synthetic data set is summarised (mean ± s.d.). Multiple recordings exist from each experimental condition (three recordings from condition 1 and two from condition 2). Here, it is difficult to judge whether the two populations differ since there is a large variation between recordings. Our aim in this paper is to develop an objective method for deciding if two populations vary where there are multiple recordings and where there is a dependence on an independent variable.

In this work, we assess the methods which were used in previous studies to assess if the correlation-distance relationship differs significantly between experimental conditions. We argue that the methods which have been previously used are insufficient given the complexity of the data and result in misleading conclusions. We propose a flexible modelling framework for data from these studies which specifies a hierarchical Bayesian model for the data which is used to assess the evidence for/against differences in correlation (we avoid the term "inference" since this is more appropriate in a frequentist framework). We demonstrate its use by applying it to data from two studies of spontaneous retinal development. We believe our framework is quite general and can be applied to a wide range of independent and dependent variables, not just distance-dependent correlations.

## Methods

In this paper we develop a new framework to test whether correlations in spontaneous retinal activity differ between two or more experimental conditions. In our case, the different experimental conditions are genotypic differences in mice (e.g. wild type versus one line of mutant mouse). Our method involves fitting a Hierarchical Bayesian model to the data which is then used to assess the evidence for/against phenotype-level differences. Most of the methods section is devoted to how we fit this model.

In this paper we use the spike time tiling coefficient (STTC) as our measure of correlation (the dependent variable) as it has been demonstrated to be highly suited to analysing correlations in spontaneous retinal activity data [15]. For brevity, we refer to the independent variable, electrode separation, as "distance". However, we believe that our method can be adapted with minimal changes to incorporate other combinations of dependent and independent variables.

## Form of the full model

For each recording (of a specified phenotype) the distribution of the correlation values at a fixed distance is modelled by a probability distribution which we refer to as the *data-generating distribution.* This distribution is assumed to be the same for all distances, recordings and phenotypes (so that all data points are parametrised identically). The parameters of this distribution (called *data-generating parameters)* depend on the distance, recording and phenotype. The data-generating parameters are specified deterministically by a function (a *distance dependence function)which* models their distance-dependence whose parameters (called *distance-dependence parameters)* depend on the recording and phenotype. To parametrise all recordings and phenotypes identically this function is assumed to be the same for all recordings and phenotypes, that is the form of the function is the same but its parameters vary. The distance-dependence parameters for each recording are assumed to be drawn from a probability distribution (a *phenotype-dependence distribution)* whose parameters depend on the phenotype *(phenotype-level parameters).*

In the Section "Modelling process" we describe how to use the data to specify the necessary distributions and functions to obtain precise form of the model.

## Alternative models

In addition to the full model described above we also evaluate three alternative models (Figure 2) to test the assumptions that there are differences between phenotypes and recordings:

- **Model A** assumes that correlations depend only on phenotype and not on recording.
- **Model B** assumes that correlations depend only on recording and not on phenotype.
- **Model C** assumes that correlations depend neither on recording nor on phenotype.
- **Model F** is the full model which assumes correlations depend both on phenotype and recording.

**Figure 2:**
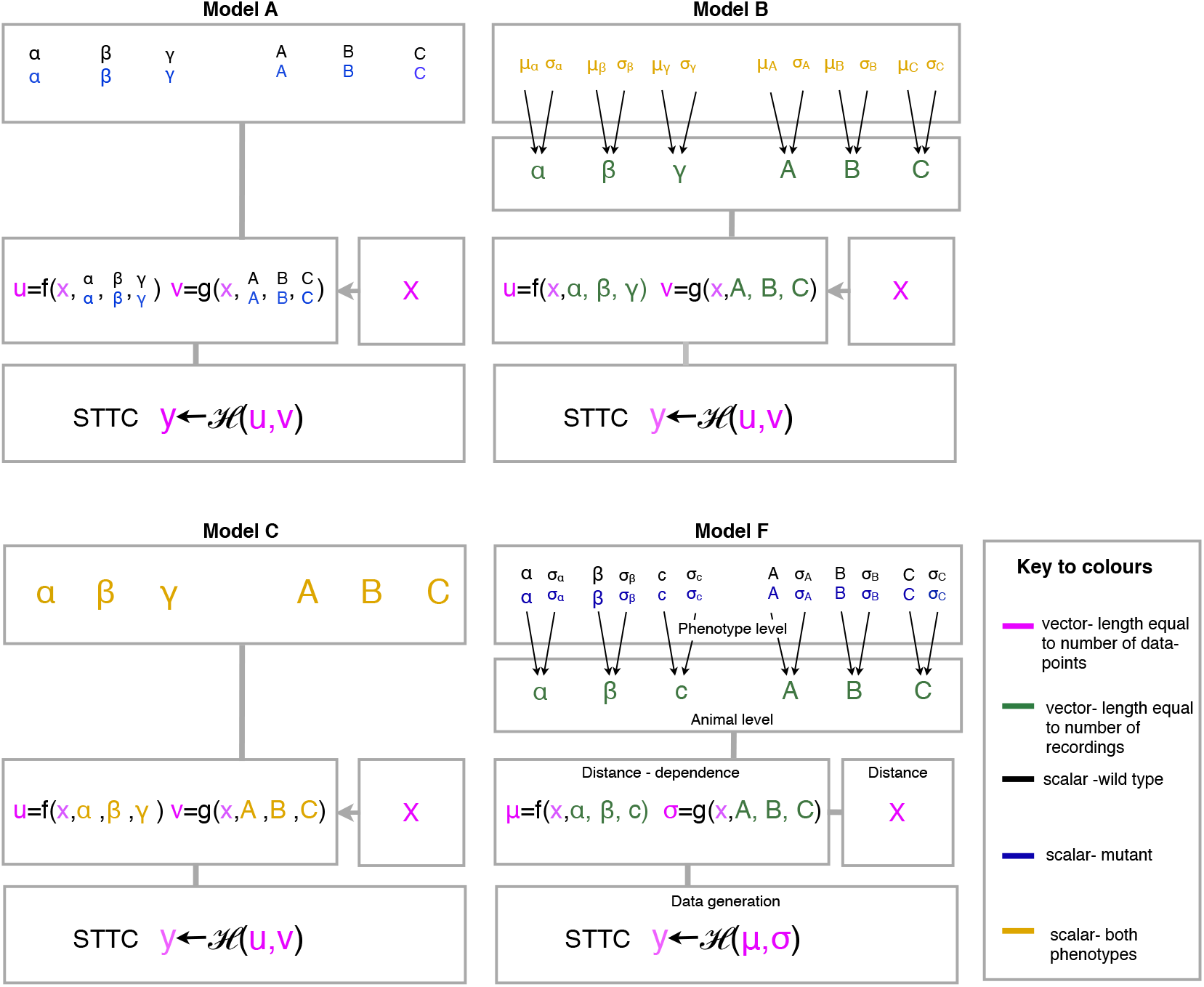
Diagrammatic representations of the four models considered. Simplifying models (A-C) are used to test importance of key assumptions in the full model (F). Model F consists of four levels. At the data-generation (bottom) level the STTC values are modelled as being drawn from a specified data-generating distribution *H* with data-generating parameters (*u, v)* which depend on the distance (*x*), the recording and the phenotype of the STTC value. At the next (distance-dependence) level the data-generating parameter values are specified deterministically by a distance-dependence function (*f, g*) which depends on the distance and which is parametrised by a set of distance-dependence parameters which depend on the recording (and in turn on the phenotype). At the next (recording) level the distance-dependence parameters for each recording are drawn from a phenotype-dependence distribution which models the variation between animals of the same phenotype. The final (phenotype) level specifies the phenotype-level parameters of the phenotype-dependence distribution. These parameters are assumed to be phenotype dependent, i.e. there is a separate value for each phenotype. Models A-C differ only in their assumptions about recording-and phenotype-level dependencies (i.e. the data-generating distribution and distance-dependence functions remain unchanged). Model A assumes no variations between recordings (i.e. distance-dependence parameters are identical for all recordings of the same phenotype). Model B assumes no phenotype-level variation (phenotype-level parameters are identical for all phenotypes). Model C assumes no recording or phenotype-level dependencies (i.e. distance-dependence parameters are identical for all recordings and phenotypes). 38

The mathematical differences in the specification of these models are described in Section "Step 3: Model recording-and phenotype-level variation".

## Modelling process

We use a hierarchical Bayesian model to investigate our data. The form of the model described previously is encapsulated in the likelihood function. To specify this, we choose well-fitting data-generating and phenotype-dependence distributions and distance-dependence functions. There is no reason to assume a specific form for these, so we use the data to find well-fitting models. The form of the phenotype-dependence distributions are chosen to be the standard model for inter-species variation (that observations are normally distributed among animals of the same species or genotype).

The process of finding the form of the likelihood function, running and testing the model and performing inference can be described in seven steps:

### Steps 1-3: specify likelihood function

**Step 1:** Choose a probability distribution to model the distribution of the correlation values at each distance. We call this the *data generating distribution.*

**Step 2:** This distribution will have associated parameters (the *data-generating parameters).* Choose functions to model the distance dependence of these parameters *(distance-dependence functions).*

**Step 3:** These functions will have associated parameters *(distance-dependence parameters).* Model their dependence on phenotype and recording (specify the *phenotype-dependence distribution* and *phenotype-level parameters).*

### Steps 4-7: run model and assess output

**Step 4:** Specify prior distributions on model parameters.

**Step 5:** Sample from the model’s posterior distributions. Plot the resulting posteriors.

**Step 6:** Assess chosen model for goodness of fit, ease of sampling and robustness. Alter the model and repeat if there are issues.

**Step 7:** Assess evidence for/against differences between phenotypes.

Figure 3 shows a schematic diagram of steps 1-3 and in the following all steps are explained in more detail.

**Figure 3:**
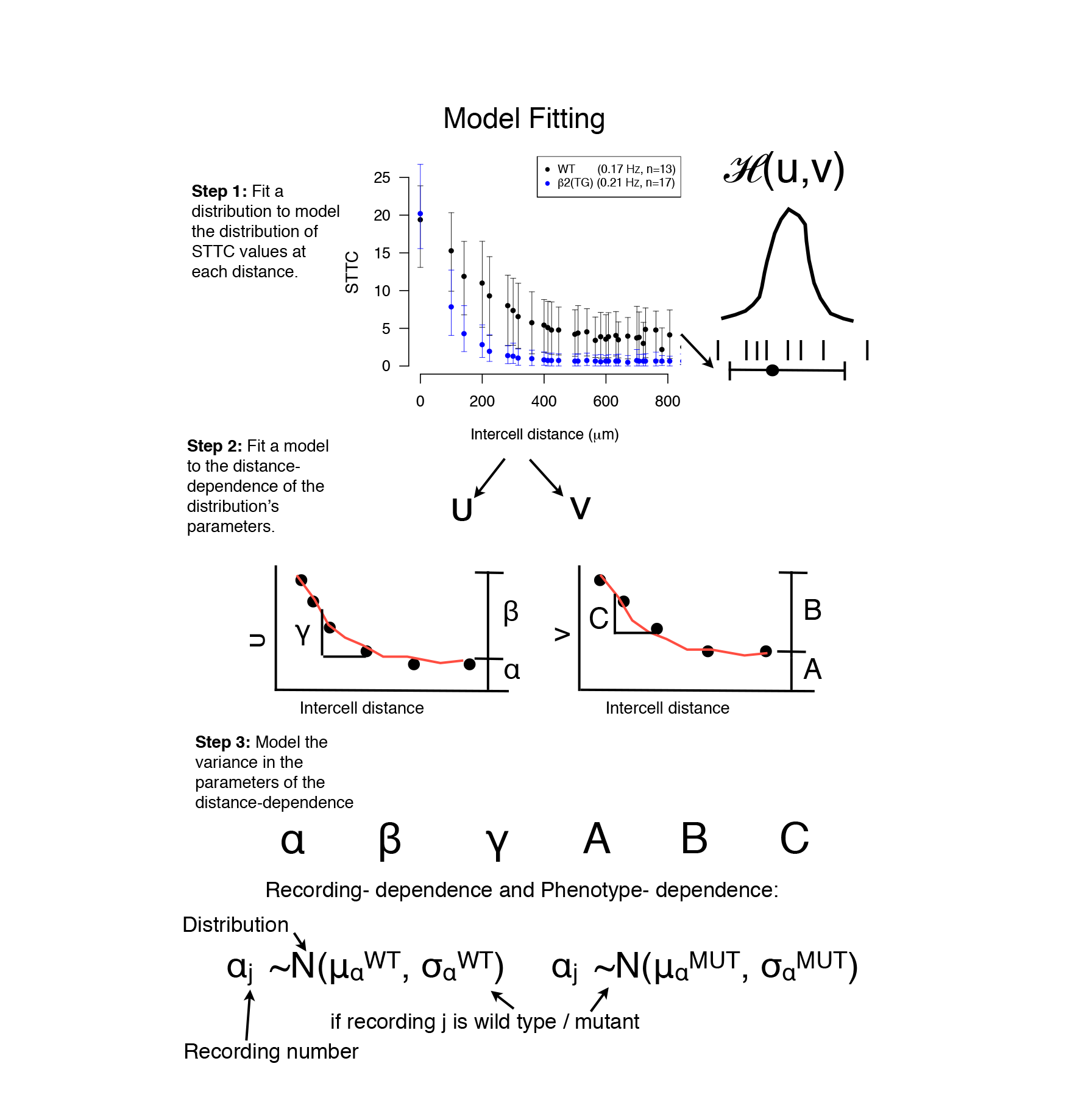
Schematic of model fitting procedure. **Step 1:** A probability distribution (*H*) is chosen to model the distribution of STTC values at a fixed distance (assumed to be the same for all distances, recordings and phenotypes). The visualisation of the mean ± standard deviation from the wild type data at 800μm is isolated and rotated. The "rug plot" (vertical lines) show the value of individual data points. Above this, the chosen probability distribution *H* is drawn. This is captured by a set of parameters (here *u* and *v*) which depend on distance, recording and phenotype. **Step 2:** we model the distance-dependence of u and v (separately) as a function (assumed identical for all recordings and phenotypes) which is parametrised by a set of parameters which have a physical interpretation. e.g. the distance-dependence of *u* can be parametrised by α, the *baseline* value of the curve, *β*, the *range* of the data: the difference between its maximum value and *α*, and *γ*, the *gradient* i.e. decay constant of the curve. The distance dependence of *u* can be modelled similarly. **Step 3** models the dependence of (*α,β,γ,A,B,C)* on recording and phenotype. Parameters for each recording are drawn from a normal distribution (the model for variation between recordings of the same phenotype) with mean and variance which are different for each phenotype (the model for phenotype-variation).

## Fitting the likelihood model: Steps 1-3

### Step 1: fitting the distance-dependence distribution

Table 1 lists the 21 continuous 1-dimensional distributions which were considered as candidates for the distance-dependence distribution and which were fitted to the data (the support of the distributions is transformed to encompass the support of the STTC if necessary). All distributions are fitted using maximum-likelihood (ML) estimates to the STTC values at each distance of each recording (i.e. not pooled). Closed-form estimates are used if they exist, if not estimates were found using a Nelder-Mead algorithm [43]. The ML estimates are graphically compared with non-parametric kernel-density estimates of the data’s distribution (using a Gaussian kernel with width set using Silverman’s rule of thumb [50]) to eliminate distributions which are poor fits. Remaining plausible distributions are ranked using the Kolmogorov statistic [35] and the squared errors between the ML estimates and the kernel-density estimates at a series of equally-spaced points. Once distributions are ranked in order of fit, their practicalities are compared: two-parameter distributions are preferred to three as this keeps model complexity to a minimum and distributions whose ML parameter estimates have a small range are preferred (this increases the chances of finding suitable starting values for Markov chain Monte Carlo (MCMC) sampling). The ranked distributions were tested for the above qualities and, if necessary, MCMC posterior samples were drawn from the candidate models to inform the decision. The importance of the trade-off meant that more sophisticated means of assessing distribution fit, e.g. the Akaike Information Criterion [2] were not needed. Several distributions were very good fits so the final choice was made on pragmatic grounds.

**Table 1.**
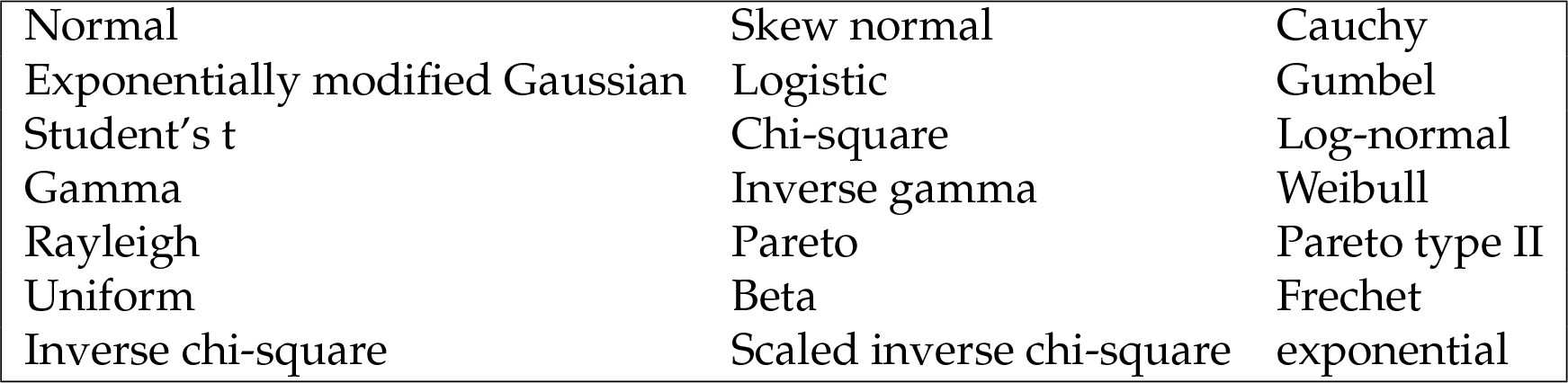
The 21 continuous one-dimensional distributions considered as data-generating distributions.

### Step 2: Fit the distance-dependence functions

The chosen data-generating distribution will have two or three (data-generating) parameters. All have a location and a scale parameter. The third parameter varies but is generally related to skewness. Plotting the ML estimates of the location and scale parameters shows a distance-dependence (concave monotonic decay with a decreasing gradient), e.g. Figure 4, which is modelled. Functions are fitted to model the distance-dependence of the scale and location parameters separately. For each parameter the same function is used for all recordings (so that parameters can be compared across phenotypes). Several candidate functions are fitted (Table 2) which have a maximum at zero and decay monotonically with increasing separation to a fixed asymptote. Exponential and reciprocal forms are considered for simplicity and because they occur frequently in the biological and physical sciences. The functions were fitted to the ML estimates of the data-generating parameters using a generalised least-squares method.

**Table 2.**
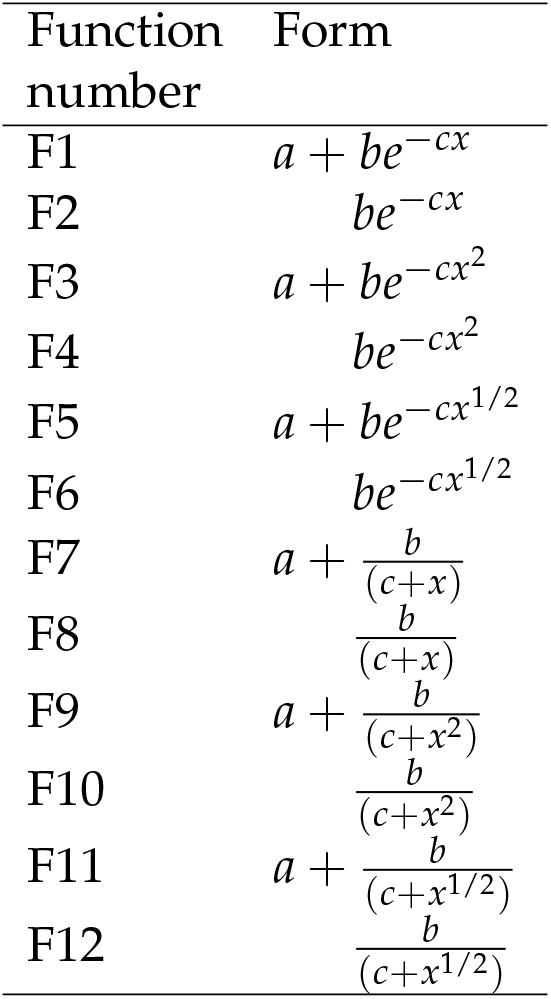
List of functions which were considered as possible distance-dependence functions of the scale and location parameters of the data-generating distribution. Functions are referred to by the identifier in column one.

| Function number | Form |
| --- | --- |
| F1 | $a + be^{-cx}$ |
| F2 | $be^{-cx}$ |
| F3 | $a + be^{-cx^2}$ |
| F4 | $be^{-cx^2}$ |
| F5 | $a + be^{-cx^{1/2}}$ |
| F6 | $be^{-cx^{1/2}}$ |
| F7 | $a + \frac{b}{(c+x)}$ |
| F8 | $\frac{b}{(c+x)}$ |
| F9 | $a + \frac{b}{(c+x^2)}$ |
| F10 | $\frac{b}{(c+x^2)}$ |
| F11 | $a + \frac{b}{(c+x^{1/2})}$ |
| F12 | $\frac{b}{(c+x^{1/2})}$ |

Functions were ranked by fit using the root mean square error (RMSE) for each recording. A trade-off is necessary between fit and practicality. Functions with fewer parameters are preferred as are those whose distance-dependence parameter fits have a small range (as this helps MCMC sampling). In Table 2 F1-6 are preferred since all parameters have direct biological interpretations: *a* - baseline value of correlation outside of waves, *b* - difference between maximum and baseline, i.e. the amount of extra correlation in waves and *c* - decay parameter, i.e. a measure of the strength of the distance-decay or alternatively, the extent of the waves. The ranked functions were assessed on their practicalities and if necessary MCMC samples were drawn to inform the decision and a function was chosen. In practice, several functions fitted the data well so the choice was made pragmatically.

When the chosen data-generating distribution had a third parameter, large variations in the parameter value were always seen and the form of the distance dependence was less clear than for the scale and location parameters. Since the quantity this parameter represents varies, the fitting procedure could not be generalised. The parameter was transformed (e.g. logarithmically) to lessen the variation and clarify its distance-dependence and appropriate functions were fitted on an ad-hoc basis. In practice the fitted distance-dependence parameters relating to the third data-generating parameter had such large ranges that sampling from their posterior distributions was impractical as it was prohibitively difficult to find appropriate starting values and convergence times were long.

### A note on Steps 1 and 2

It is more commson to fit both the data-generating distribution (sometimes called the "error distribution") and the distance-dependence function at the same time as this ensures that the best fitting model is chosen. This was impractical for the data sets chosen and the large number of candidate data-generating distributions and distance-dependence functions, so the two stages were performed separately.

### Step 3: Model recording-and phenotype-level variation

The chosen distance-dependence function has a set of distance-dependence parameters whose dependence on recording and phenotype are modelled. To illustrate this, suppose the Gumbel distribution [26] is the data-generating distribution and the distance-dependence functions for the location and scale parameters (any third parameter is considered identically) are F1 and F3 respectively. Then

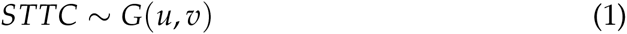

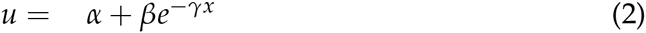

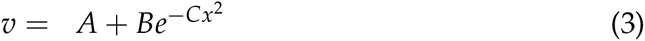

where *x* is the electrode separation.

The distance-dependence parameters are (*α, β, γ*) which dictate the value of the data-generating location parameter and (*A, B, C*) which dictate the data generating scale parameter (we use this convention throughout). In the full model (F) the parameter-value for each recording is assumed to be normally-distributed with a phenotype-dependent mean and standard deviation. We call these parameters the phenotype-level mean and phenotype-level standard deviation.

For instance, if a recording *j* is of phenotype Z (denoted *j*(Z)) then its *α_j(z)_* distance-dependence parameter is modelled as:

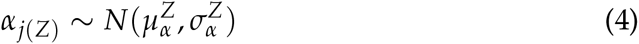

Where 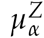 and 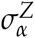 are the phenotype-level parameters. Likewise if recording *i* is of phenotype ϒ then its parameter *α_i_ (ϒ)* is drawn from a Normal distribution with parameters 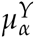 and 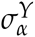.

Models A, B and C vary these assumptions. Model A assumes no recording-level differences i.e. 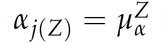 for all recordings of phenotype Z and 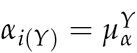 for all recordings of phenotype Y. Model B assumes no phenotype-differences i.e. 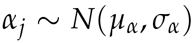 regardless of phenotype. Model C assumes no recording-or phenotype-differences i.e. *α_j_ ∼ N(μ_α_, σ_α_)* regardless of phenotype. Model C assumes no recording-or phenotype-differences i.e. *α_j_ = μ_α_* for all phenotypes.

### Step 4: Specifying the priors

The first three steps generate the likelihood function; priors on the phenotype-level parameters are needed to complete the Bayesian model.

There may be constraints on the phenotype-level parameters which are specified by the likelihood function. We have no extra knowledge about them, so we choose an uninformative prior. The size of the model and data sets meant that it was impractical to perform MCMC sampling without specifying bounds on the parameters. It was therefore efficient to choose maximum entropy prior [12] as our uninformative prior, as for a bounded, continuous support, it is the uniform distribution [30]. In practice, many parameters are naturally bounded by the range of the STTC and the forms of the likelihood function. For those which were not, conservative bounds were chosen (see Results).

### Step 5: Running the model

Models were implemented in Stan [56] and convergence was determined using the in-built Gelman-Rubin [23] diagnostic test (with values 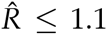 being adequately converged [23]) and verified by inspecting trace plots of the samples. Six separate simulations each generating 15,000 samples were run for each model to ensure robustness [6]. Depending on the model, convergence required 16,000-50,000 iterations and took between 24 and 100 hours. Twice the number of iterations required to achieve convergence were used as the burn-in period, after which, samples were saved. All posterior distributions and synthetic data shown in Figures were taken from/generated from six merged MCMC chains each of 15,000 samples.

### Step 6: Model assessment

Plots and summary statistics (deciles) of the posterior distributions were compared between simulations to ensure all runs converged to the same distributions.

Posterior predicative sampling [49] is used to assess the fit of the model: 2000 repeats of draws from the posterior distribution of the model (for each data point 2,000 synthetic data points are generated from the model using the distance which corresponds to the data point and sampling from the posterior distributions which match the phenotype and recording of that point) and summary statistics are compared between data and the synthetic data for each recording. (One minor issue is that in principle our simulated data can generate values outside the bounds of possible values [−1,+1] for STTC. However, in practice, this happens rarely, and so we simply discard those values.) The median and interquartile ranges are compared since these are not parametrised by the chosen data-generating distribution (Gumbel) but are close in interpretation to its scale and location parameters.

The assumption that the phenotype-dependence distribution is normal was assessed using pivotal density measures [31, 70]. For each parameter (e.g. *α*), a draw was made from its posterior distribution from each recording *(α_j_)* and a draw was made from the posterior distributions of the corresponding phenotype level mean 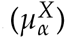 and standard deviation 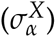 where X is the phenotype of recording *j.* Then

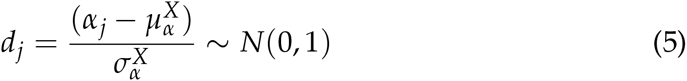

under the model and is therefore a pivotal density measure [31] and

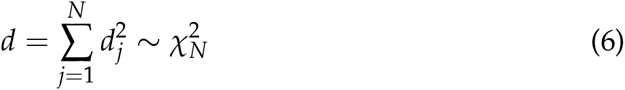

where N is the number of recordings. 2,000 replicates of d were generated and the distribution compared to the theoretical distribution (formalised tests exist [70] but were found to be hyper-sensitive).

The robustness of the conclusions to perturbations in the model’s assumptions was also tested. Altering the distance-dependence functions was not possible as this would alter the parameters so a different inference was being made. Instead, model robustness to the choice of data-generating distribution and prior is tested by perturbing these distributions and testing if they affected the resulting inference. The data-generating distribution was perturbed (from Gumbel) to normal and the prior is perturbed (from uniform) to normal with a small standard deviation and a mean which is off-set from the range of the posterior which is obtained using a uniform prior. These perturbed models are run and assessed as described previously. The perturbations are large (as opposed to subtler perturbation methods such as mixture distributions/priors) as it is important that the conclusions are robust and remain unchanged under different but still plausible models as this means they are more likely to reflect something inherent about the system as opposed to being an idiosyncrasy of one model which happens to fit the data well.

The Watanabe-Akaike information criterion (WAIC) [64] is used to assess the relative impact of the assumptions made about recording-and phenotype-level parameters of the model. This criterion accounts for model complexity and is used to compare the fits of Models A, B C and F.

### Step 7: Assessing evidence for/against differences between phenotypes

Evidence for the existence of phenotype-level differences is obtained by comparing the WAIC values of Model F and Model B (which does not model phenotype-differences but which is otherwise identical to F). If the WAIC value of Model F is sufficiently lower than that of Model B to provide evidence in favour of Model F (Δ*WAIC* ≥ 5 according to general guidelines [11]) then this is evidence that there are phenotype-level differences in the data. If this is not the case we either conclude that both models are equally parsimonious or that Model B is preferred and there is no evidence for phenotype-level differences.

In addition to this, we compare the posterior distributions of the phenotype-level parameters to ascertain in which features the correlations differ (and to provide further evidence for/against differences between phenotypes). We use the following ad-hoc test: if the 95% highest posterior density (HPD) regions are disjoint between phenotypes this is evidence that there are differences between phenotypes in this feature (parameter).

### Presentation of data

In Figures 4, 5, 8 and 11 we use box plots to summarise the data for each phenotype (pooled across all recordings of the same phenotype). In all box plots the "box" consists of the median and first and third quartiles of our data. The "whiskers" show the largest and smallest values which are 1.5 times the interquartile range outside the box. Points which fall outside the whiskers are plotted individually as outliers.

## Data sets used

We apply the framework described above to two different data sets to demonstrate its use.

### Data from Xu et al. [69]

This paper compared the spontaneous retinal activity and retinotopic and eye-segregation maps of wild type mouse and a mutant genotype *β2(TG)*. A correlation-distance plot was key to the analysis and demonstrated that the *β*2(TG) mutant has truncated waves compared to wild type (that is, its correlations decay much more strongly with distance). We apply the method described previously to this data set to determine if the correlation-distance profiles differ between the two phenotypes. The data set is summarised in Figure 4.

**Figure 4:**
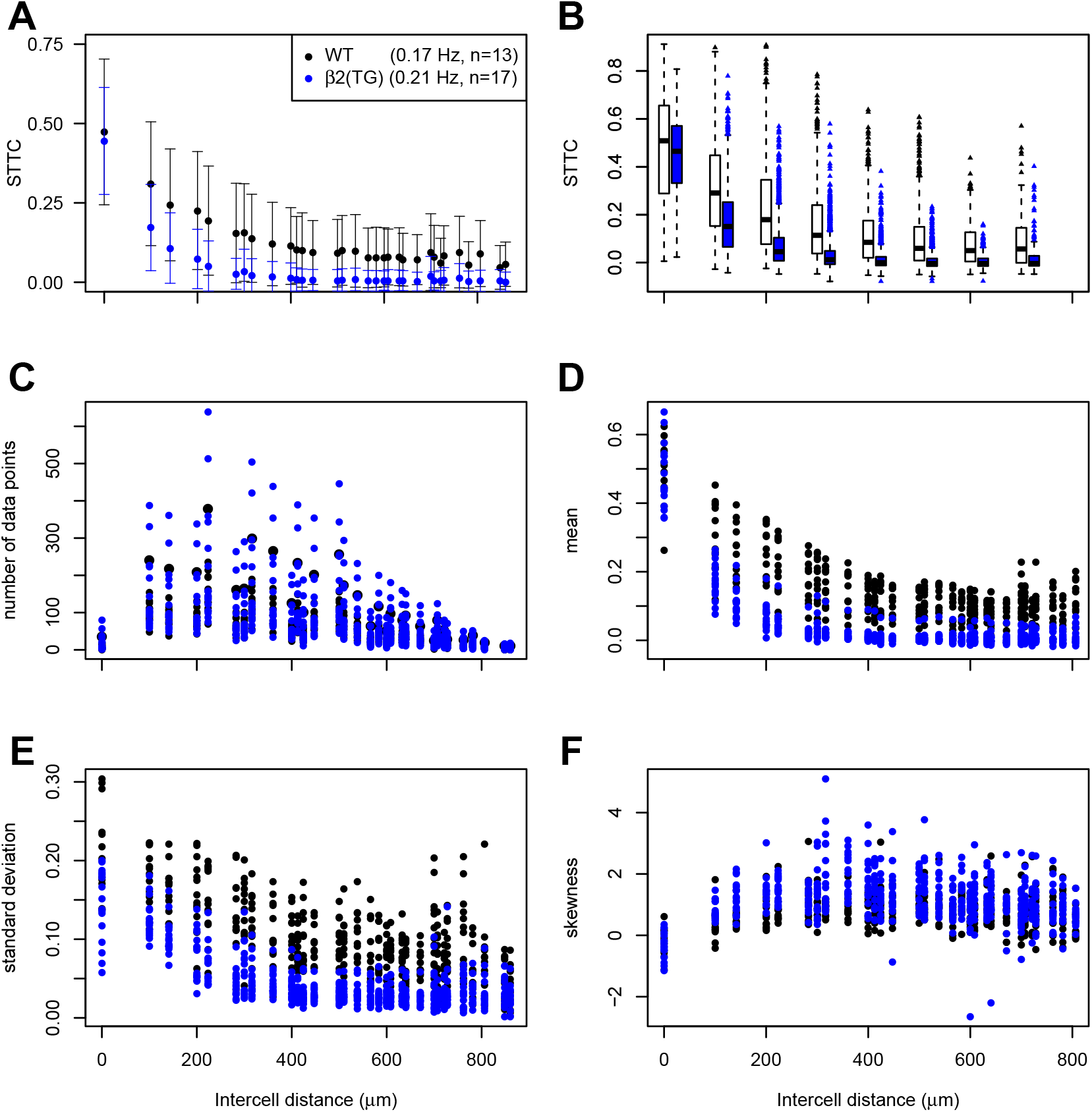
Exploratory analysis of STTC data from Xu et al. [69].A: Mean and ± standard deviation of STTC values pooled across recordings by phenotype at each recorded distance. B: Boxplots (as described in Methods) of the wild type and *β*2(TG) STTC values pooled across recordings by phenotype at a subset of all possible distances (every 100 μm). C: The number of STTC values at each distance for all recordings (each recording contributes one point at each distance). D: Mean values of STTC at each distance for all recordings. E: Standard deviation of STTC values at each distance by recording. F: Skewness of STTC values at each distance by recording.

The data from this paper is freely available from the CARMEN project [21] and consists of 30 MEA recordings of spontaneous retinal activity, 13 wild type post-natal day (P) 4 and 17 *β2(TG)* (P4). Most recordings are over 1,200 seconds. The spike times are pre-sorted and each recording contains between 40 and 118 separate spike trains.

### Data from Blankenship et al. [5]

This paper compared the spontaneous retinal activity and eye-specific segregation in the lateral geniculate nucleus in wild type mouse and two mutant genotypes Cx45ko and Cx36/45dko which lack either one or two neuronal connexins. A correlation-distance plot was used to demonstrate the differences in correlations. This data was later re-analysed using the STTC [15] using a correlation-distance plot (Figure 5A) from which it was concluded that the correlations do not differ greatly between wild type and Cx45ko, but Cx36/45dko has lower correlations and weaker distance-dependence than the other two phenotypes. The data set is summarised in Figure 4.

The data is freely available as before [21] and consists of five wild type, four Cx45ko and six Cx36/45dko recordings. Recordings range in duration from 3,130-6,270 seconds and each has between 47 and 111 spike trains.

**Figure 5:**
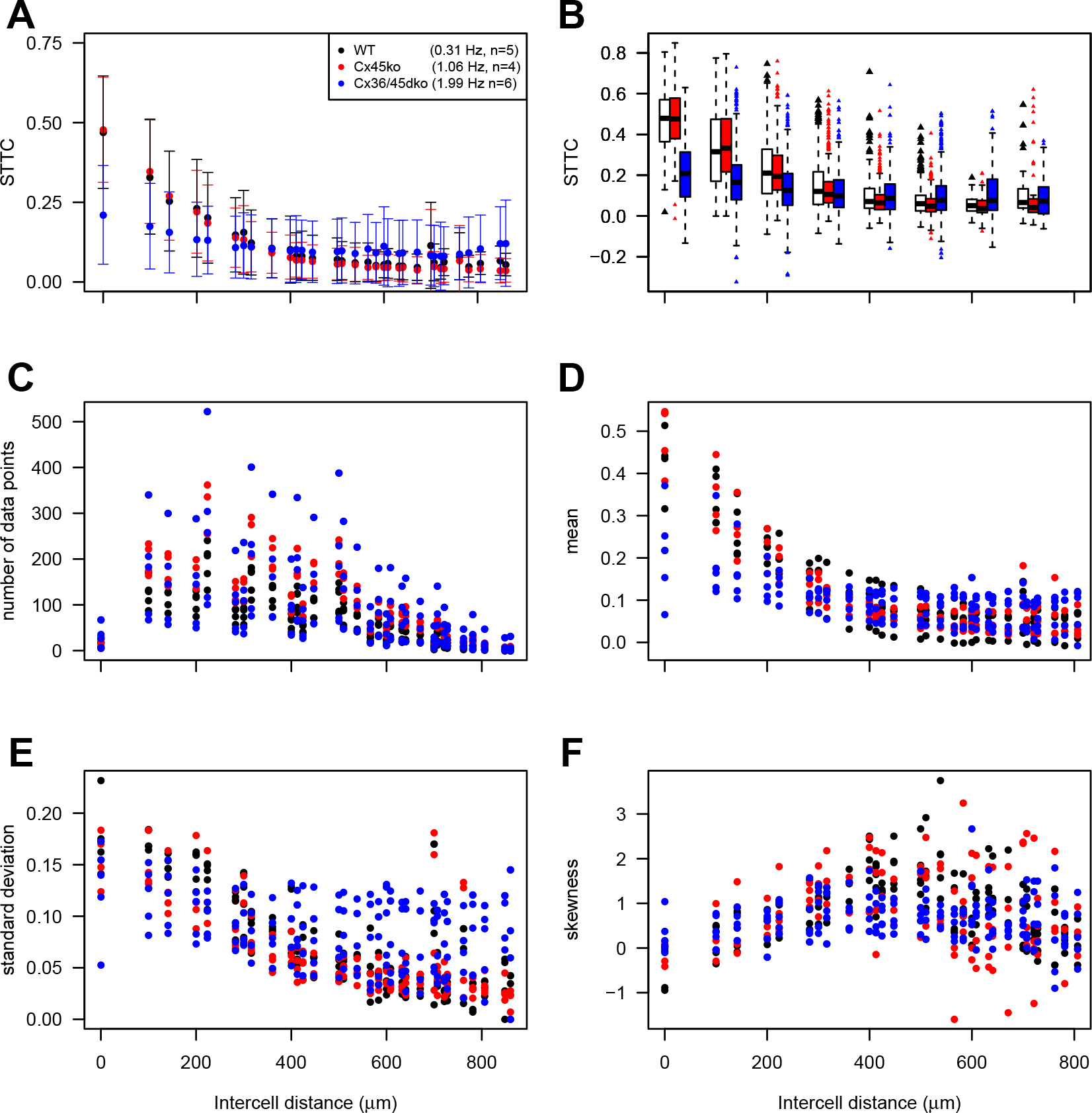
Exploratory analysis of STTC data from Blankenship et al. [5].A: Mean and ± standard deviation of STTC values pooled across recordings by phenotype at each recorded distance. B: Boxplots (as described in Methods) of the wild type, Cx45 ko and Cx36/45dko STTC values pooled across recordings by phenotype at a subset of all possible distances (every 100 μm). C: The number of STTC values at each distance for all recordings (each recording contributes one point at each distance). D: Mean values of the STTC at each distance for all recordings. E: Standard deviation of STTC values at each distance by recording. F: Skewness of STTC values at each distance by recording.

There are three phenotypes in this data set so we can test for evidence of differences between phenotypes in general and between two phenotypes specifically. We introduce a new Model G therefore to test for evidence for/against differences between wild type and Cx45ko. This model is identical to Model F except that the wild type and Cx45ko phenotypes are considered to be identical so are described by the same phenotype-level parameters and posterior distributions. The Cx36/45dko phenotype is considered distinct and modelled separately. Then, if Model G is preferred to Model F (according to the WAIC) this is evidence that there are no significant differences between wild type and Cx45ko.

## Results

Distance-dependence of correlations in neural activity are typically assessed by calculating pair-wise correlations from MEA recordings and plotting correlation as a function of the separation of the electrodes from which the neutrons were recorded [66]. Since the distance dependence of these correlations is thought to be instructive in topographic map formation [65], it is common to use these plots to compare the distance-dependence of correlations between different phenotypes/experimental conditions. Visual inspection of these plots is, in itself, not sufficient to draw robust conclusions as to whether there are differences between phenotypes. It is necessary to statistically assess the evidence for (or against) differences in correlations between phenotypes.

### Previously used methods of inference are inadequate

Table 3 lists all studies which compared the correlations of two or more experimental conditions using the correlation index. Many of them include a correlation-distance plot, although several plot and compare correlations at only one distance. The method of inference used in each study is noted, as is the method used to model the distance-dependence of correlation (if included). Over half the studies did not use any formalised method of inference and simply appealed to inspection of the correlation-distance plots to support their conclusion. The remaining studies used standardised significance tests which can be group according to whether they compare two conditions (student’s t-test, Mann-Whitney U test, Mood’s Median test) or two or more conditions (oneway ANOVA and Kruskal-Wallis ANOVA) and whether they assume that the data is normally distributed (student’s t-test, one-way ANOVA) or do not make this assumption (Mann-Whitney U test, Mood’s Median test and Kruskal-Wallis ANOVA).

**Table 3.** Publications using the correlation index to compare conditions grouped by method of inference. Bold headings denote the type of significance testing used. The publication is listed in column 1; column 2 notes the method, if any, for modelling the distance-dependence of the correlation.

| **Inference type** | **Modelling** |
| --- | --- |
| **No inference** |  |
| Wong et al. [66] | Linear regression on log CI |
| Butts and Rokhsar [8] |  |
| Demas et al. [16] | Exponential fit<br>+ 95% confidence intervals |
| Cang et al. [10] |  |
| Bjartmar et al. [3] |  |
| Demas et al. [18] |  |
| Sun et al. [59] |  |
| Godfrey et al. [25] |  |
| Godfrey and Eglen [24] | Exponential fit |
| Stafford et al. [55] |  |
| Hennig et al. [27] |  |
| Ding et al. [20] |  |
| Xu et al. [68] |  |
| Xu et al. [69] |  |
| Speer et al. [53] |  |
| Blankenship et al. [5] |  |
| Blank et al. [4] |  |
| Chabrol et al. [13] |  |
| Dhande et al. [19] |  |
| Soto et al. [52] |  |
| Demas et al. [17] | Exponential fit |
| Kirkby and Feller [33] |  |
| Cain et al. [9] |  |
| Eglen et al. [21] |  |
| Lee et al. [36] |  |
| Maccione et al. [38] | Exponential fit |
| **Student's t-test** |  |
| McLaughlin et al. [42] |  |
| Wang et al. [63] |  |
| Huang et al. [28] |  |
| Speer et al. [54] |  |
| Xu et al. [67] |  |
| **Mann-Whitney U test** |  |
| Stasheff [57] |  |
| Sun et al. [60] |  |
| Zhang et al. [71] |  |
| Stasheff et al. [58] |  |
| **Mood's Median Test** |  |
| Torborg et al. [62] |  |
| **One way ANOVA +**<br>**Fisher's PLSD Post-hoc test** |  |
| MacLaren et al. [39] |  |
| **Kruskal-Wallis ANOVA +**<br>**Dunn's Post-hoc test** |  |
| Personius and Balice-Gordon [45] |  |
| Torborg et al. [61] |  |
| Personius et al. [46] |  |
| Personius et al. [47] |  |
| Chiang et al. [14] |  |

The common feature of these tests is that they only compare the distribution of the data at one fixed distance and determine whether correlations differ at that difference. Some studies (e.g. [57]) performed this test at one difference (usually at zero electrode separation) and others (e.g. [60]) performed them at all distances.

Given that the distance-dependence of the correlations is the feature which is implicated in map formation, it is surprising how few studies attempted to model it. Those that did fitted an exponential decay but did not use this in the inference. Performing significance tests at one distance or at each distance separately does not capture the distance-dependence and loses vital information about the data.

This loss of information is concerning since the differences in the correlation-distance graphs are often key to the overall conclusion of the paper. A further piece of information in the data is the variance which is seen between recordings of the same phenotype (which can be large, see Figures 4 and 5) is lost as the data is pooled across recordings at each distance. Since both these factors are likely to impact the inference, a credible inference must account for both. A further limitation of the current approaches is that they cannot distinguish how correlations differ: a method which can do this could highlight specific features of the correlation-distance relationship which may be implicated in map development.

### Modelling approach

Since the previous methods used to compare correlations between phenotypes are problematic, we aim to develop a framework which allows us to assess evidence for/against differences in the correlation-distance relationship between phenotypes in a credible manner.

We require that such a framework be robust and intuitive. As we wish that it to be used on MEA data to replace the current standard statistical tests, it is important that the method we propose requires minimal increases in terms of conceptual understanding and computational effort. Robustness is important to ensure that any effect measured represents an underlying biological phenomenon and not an idiosyncrasy of a model which happens to fit the data well.

We use a Bayesian approach to modelling partly due to the high-profile criticisms of the use of standard frequentist tests in the biological sciences in general [29] and neuroscience in particular [7], and also because Bayesian modelling allows us to incorporate all relevant information (e.g. inter-recording variance) in a straight-forward and flexible way. An additional benefit of using Bayesian analysis in the biological sciences is that relevant information (e.g. from other studies) can be incorporated into models in the form of the prior distributions.

Previous methods of inference have ignored the distance-dependence of the correlations and the inter-recording variation, both of which are likely to be key variables which explain the variations in the data. We therefore fit a Bayesian, hierarchical model as described in Methods. As far as we are aware, there is no biological evidence which suggests that the model should take a certain form (e.g. that correlation values at one distance should be assumed to be normally distributed or that the distance-dependence is exponential). We therefore fit a variety of possible distributions and functions of data and compare fits to choose a final model. In this sense our model is data-driven as opposed to hypothesis-driven.

The form of the model which we use and the steps taken to fit, run and assess the model and to evaluate the evidence from the model for/against differences in phenotypes are presented in Methods. To avoid repetition, the rationale for the model is briefly noted in Methods.

## Modelling data from Xu et al. [69]

### Modelling process

The modelling process and the data set have been described in Methods. The best-fitting data-distribution was found to be the Gumbel distribution which has a location parameter u and a scale parameter v and the distance-dependence functions F1 and F3 were the best fits to model the distance dependence of u and v respectively (Figure 6). The full model is therefore:

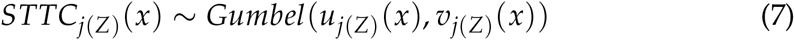

where *x* denotes the electrode separation at which the correlation value is measured, *j* is the recording number and *j(z)* denotes that recording is from phe-notype Z (here the two phenotypes are wild type and *β2(TG))*. *u* and *v* are the parameters of the Gumbel distribution which depend on the recording (and therefore the phenotype) and the distance.

**Figure 6:**
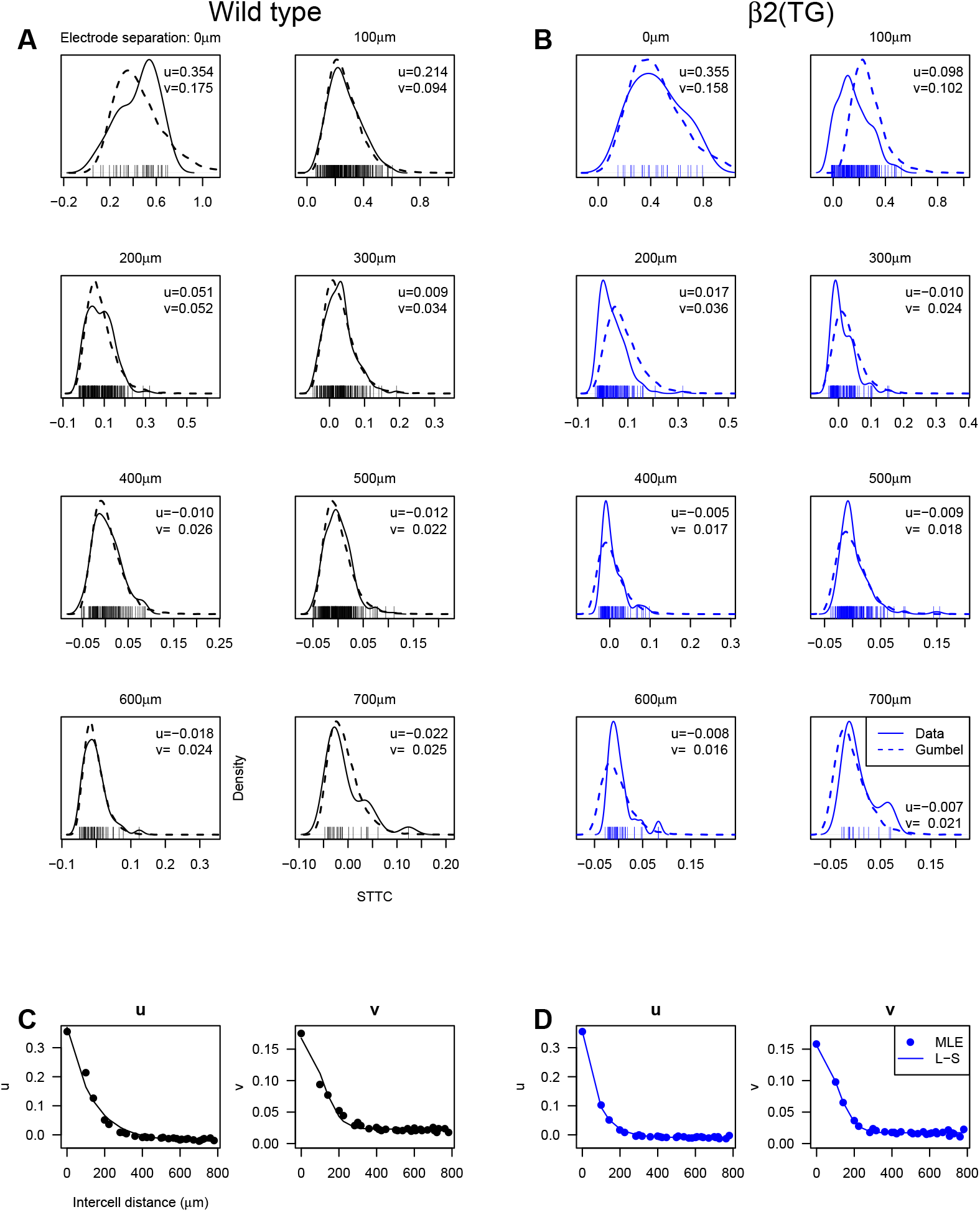
Maximum likelihood (ML) fits of the data-generating distribution and least-squares (LS) fits of the distance-dependence functions to data from Xu et al. [69]. A: Non-parametric kernel density estimates (solid lines) of the data-generating distribution at a subset of all recorded distances (every 100 μm) in one wild type recording. Rug plots at bottom show the STTC values being smoothed. The ML fits of the chosen data-generating distribution (Gumbel) are shown as dotted lines, and the ML estimates of the distribution’s parameters (*u* and *v*) are written in the top-right corner. B: One *β*2 (TG) recording shown in same format as A. C,D: Plots of the ML fits of the location *u* and scale *v* parameters of the Gumbel distribution (points) at all recorded distances for the wild type recording used in A (C) or B (D). The least-squares fits of the distance-dependence functions (F1 for u, F3 for v; Table 2) are shown as lines.

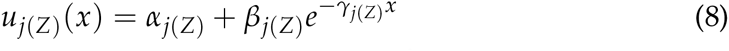

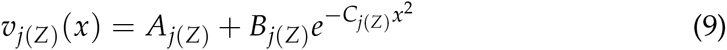

The six recording level parameters 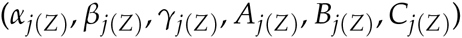 all have identical dependence on the phenotype:

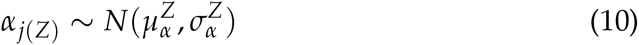

Where *N* is the normal distribution. For brevity we drop the phenotype indicator from the phenotype-level parameters (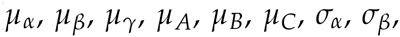 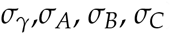), leaving it understood that their values depend on the phenotype and use color to distinguish between them in the figures.

The finite range of the STTC ([–1,1]) and the form of the distance dependence-functions allows us to put bounds on the ranges of the phenotype-level parameters. The range of the STTC implies –1 ≤ *u* ≤ 1 and from this we deduce –1 ≤ *μ*_α_ ≤ 1 and 0 ≤ *μ_β_* ≤ 2. *μ_γ_* must be positive (since u decays with distance) and since the scale of decay is hundreds of micrometers, 0 ≤ *μ_γ_* ≤ 1 is a conservative bound. Bounds can be placed on the mean parameters of *v (μ_A_, μ_B_, μ_C_)* and the standard deviation parameters 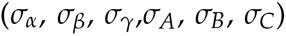 using Popovicius inequality [48] on variances which bounds the variance of any bounded probability distribution.

This bounds the variance of the STTC:

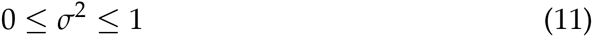

Variance is related to the Gumbel distribution’s scale parameter *v* by the following [26]:

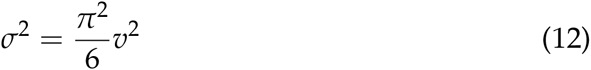

From which the following constraints follow: 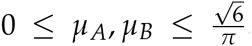. *μ_c_* is constrained using the same argument used to constrain *μ_γ_* so 0 ≤ *μ_C_* ≤ 1. The standard deviation parameters are constrained similarly using Popoviciu’s inequality.

MCMC samples from the model are generated as described in Methods using uniform priors over the bounds of the phenotype-level parameters.

### Model assessment

The results of the modelling process are the posterior distributions of all the model’s parameters. Since we are interested in phenotype-level differences we only present the posterior distributions of the phenotype-level parameters (Figure 7).

**Figure 7:**
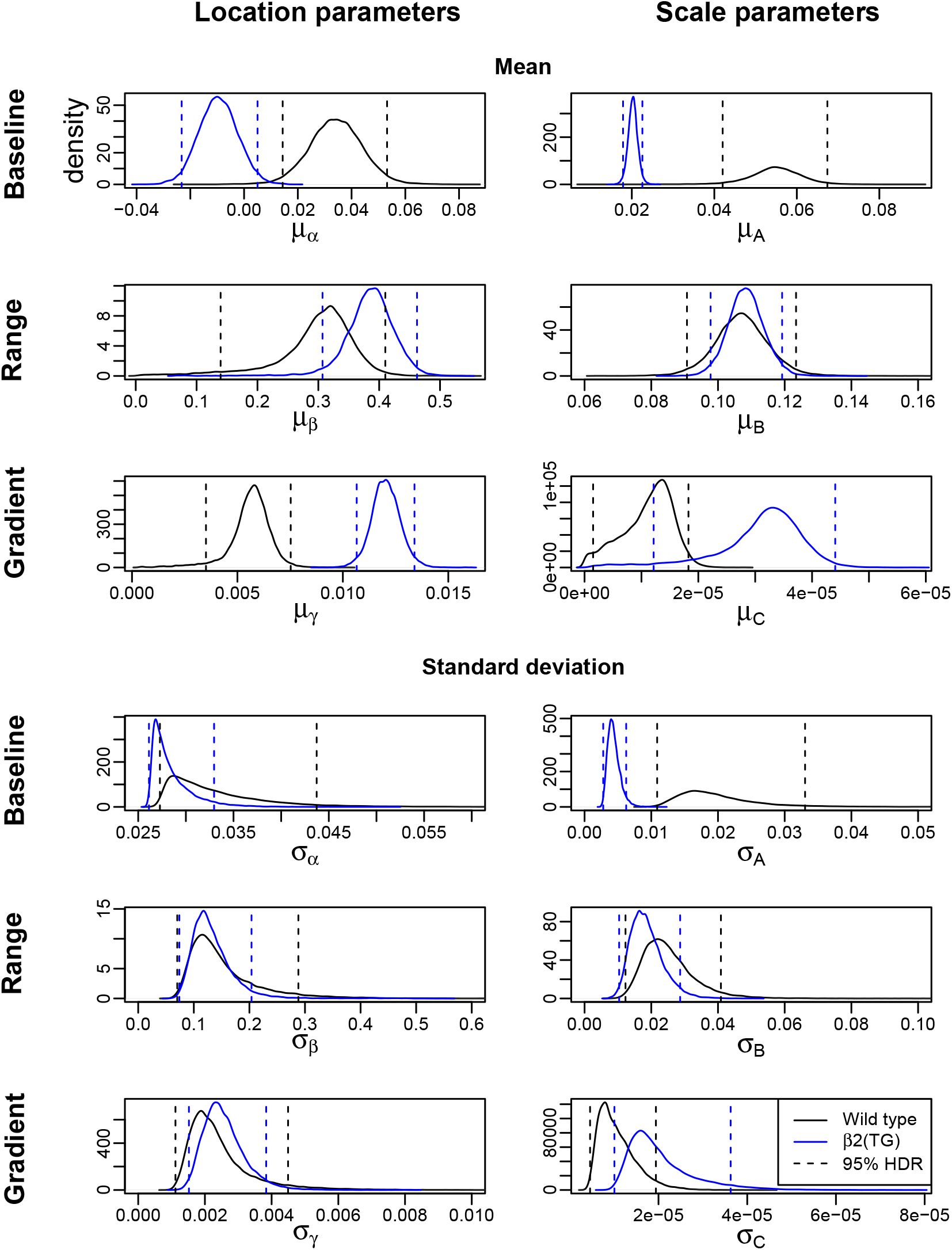
Posterior distributions of the phenotype-level parameters for Model F with Xu et al. [69] data. Parameters are modelled separately for each phenotype and the posterior distributions of each are plotted. Parameters are grouped according to whether they relate to location (*u*) or scale (*v*) parameter of the Gumbel distribution and if they set the mean or the variance across recordings. The biological interpretation of each parameter (which feature of the correlation-distance graph it relates to) is noted at the side. Posterior distributions were generated as described in Methods (Step 5). The dotted lines denote the 95% highest density region for each curve.

Assessment of the model’s ability to match the data (Figure 8A and B) demonstrates that the model is able to match the data well. The main cause of discrepancy between the model’s synthetic data and the data is that the stochastic nature of the model means that it can produce synthetic data with values ≥ 1 which is not biologically feasible as these values fall outside the range of the STTC [15].

**Figure 8:**
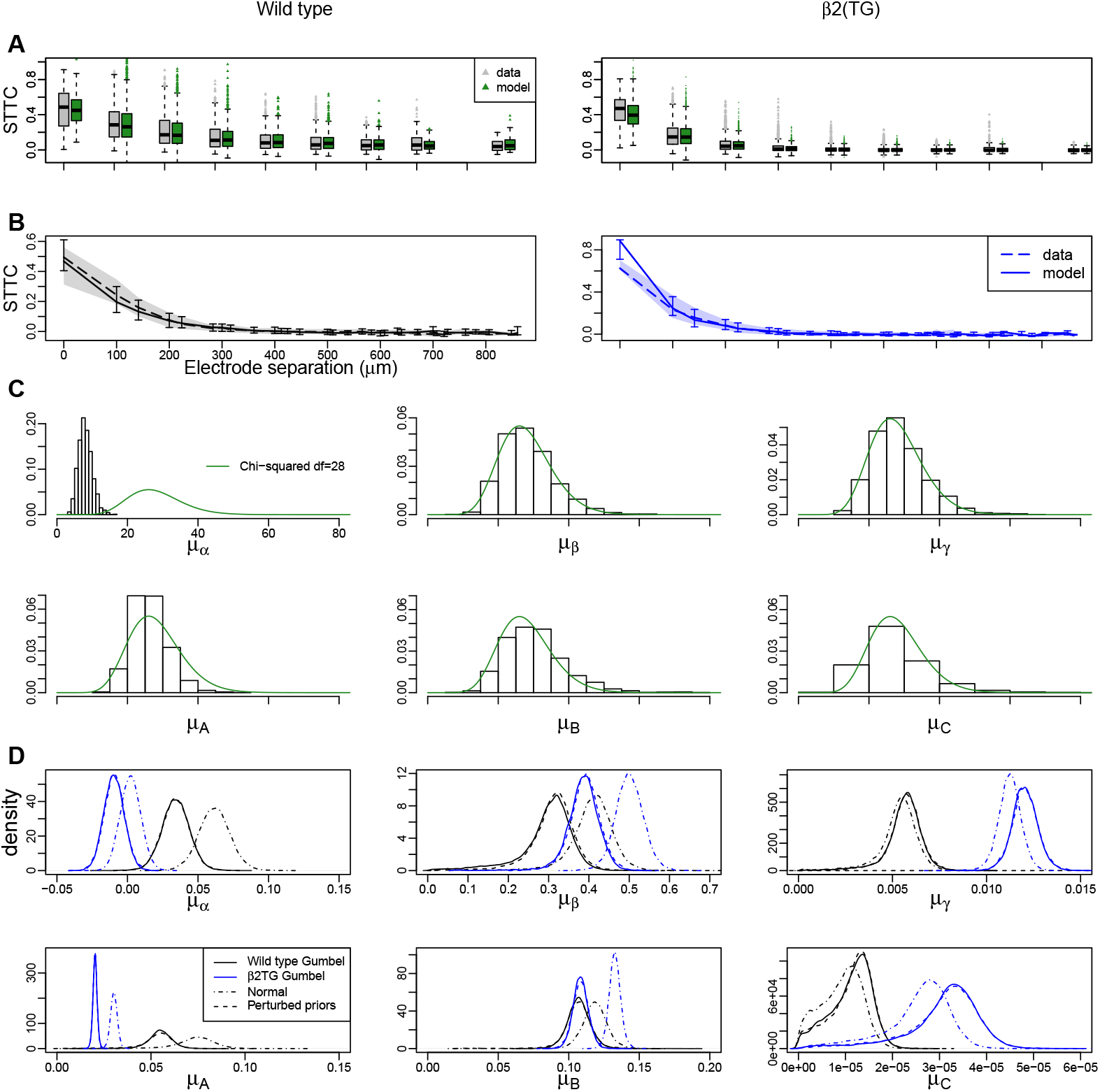
Assessment of Model F of data from Xu et al. [69]. A: Box plots comparing data and synthetic data generated using posterior predicative sampling. Data (recorded and synthetic) is pooled across all recordings by phenotype (left: wild type, right: *β2(TG))*. A sub-set of all recorded distances is shown (every 100 μm). B: Median and interquartile range of data (indicated by solid line and bars) from two recordings (left: wild type, right: *β2 (TG))*and synthetic data (indicated by dotted lines and shading). In Panels A and B for each recorded STTC value, a synthetic data point is generated with the same phenotype, recording and distance. C: Pivotal density measures (PDM; see methods, Step 6) for each phenotype-level mean parameter. Histograms of 2,000 PDM replicates are plotted along with the theoretical distribution *(x^2^* with 28 degrees of freedom; two recordings were removed as outliers). D: The posterior distributions of the phenotype-level mean parameters (Methods, Step 5) of Model F are plotted along with posterior distributions from two perturbed models where either data-generation distribution was assumed to be normal, or where the prior distributions were assumed to be normally distributed *[μ_α_ ∼ N*(0,0.5), *μ_β_ ∼ N*(1,0.5), *μ_γ_*, *μ_C_* ∼ *N*(0.25,0.25), *μ_A_*, *μ_B_* ∼ *N*(0.5,0.5)].

Assessment of the model’s performance at the recording level (Figure 8C) shows that the assumption of a normal distribution is sound. *μ_α_* is slightly under-dispersed compared to the theoretical distribution but this is not a concern as there is no reason to suppose the generation of this parameter differs from that of any other and the model fits the data well.

Assessment of the model’s robustness to both its prior and its assumptions (Figure 8D) show that the conclusions do not change. The perturbation to the prior used is strong and yet the posterior distributions produced are almost identical to those generated using a uniform prior. This implies that the data dominates the prior which does not have undue effect. The perturbation to the data-generating distribution shown in Figure 8D is also strong (from a Gumbel to a Normal distribution). This results in shifts in the posterior distributions (reflecting the different parametrisations) however, the inference will not change since posterior distributions which are disjoint for each phenotype remain disjoint and those which overlap, still overlap in the perturbed form.

**Table 4.** WAIC values for each model, listed in increasing order, of the Xu et al. data [69].

| Model | WAIC |
| --- | --- |
| F | -197999.3 |
| B | -197994.2 |
| A | -183034.5 |
| C | -148433.7 |

Assessment of the importance of including phenotype-level differences and recording-level differences on model fit is performed using the WAIC (Table 4). Model F is the most parsimonious (best fitting accounting for complexity) closely followed by the model (B) which has recording differences, but no phenotype level differences. The difference in WAIC between Models F and B (∼ 5.1) is sufficiently large to provide evidence to support the use of Model F over Model B according to generally-held guidelines [11]. There is a large improvement in fit between Model B (recording-differences only) and Model A (phenotype-differences) only which highlights the importance of including inter-recording variation in modelling.

**Table 5.** Widths of 95% highest posterior density (HPD) regions for Models A-F on Xu et al. data. The HPD widths of all phenotype-level mean parameters are shown for each model. For models A and F where the parameter is phenotype-level dependent, the HPD widths for both wild type and *β2(TG)* are given. Models B and F include a recording-dependence and appear in bold. Narrower HPD regions indicate higher confidence in the location of the parameters. Widths are given to one significant figure, sufficient to compare widths among models.

| Parameter |  | Model A | Model B | Model C | Model F |
| --- | --- | --- | --- | --- | --- |
| $\mu_\alpha$ | WT | 0.005 | **0.02** | 0.001 | **0.04** |
| | $\beta 2(TG)$ | 0.0007 | | | **0.03** |
| $\mu_\beta$ | WT | 0.03 | **0.1** | 0.02 | **0.3** |
| | $\beta 2(TG)$ | 0.02 | | | **0.2** |
| $\mu_\gamma$ | WT | 0.0007 | **0.04** | 0.0004 | **0.004** |
| | $\beta 2(TG)$ | 0.0005 | | | **0.003** |
| $\mu_A$ | WT | 0.003 | **0.04** | 0.001 | **0.03** |
| | $\beta 2(TG)$ | 0.0005 | | | **0.005** |
| $\mu_B$ | WT | 0.01 | **0.02** | 0.006 | **0.03** |
| | $\beta 2(TG)$ | 0.007 | | | **0.02** |
| $\mu_C$ | WT | 0.000002 | **0.00003** | 0.000001 | **0.00002** |
| | $\beta 2(TG)$ | 0.000002 | | | **0.00003** |

The posterior distributions of parameters from models which have no recording-level differences (models A and C) are much more localised than those which include recording differences (Table 5), making them over-confident in the parameters’ locations (as has been noted in frequentist statistics [1]). This further supports the importance of modelling inter-recording differences.

### Assessing evidence for/against phenotype-level differences

The difference in WAIC values between the full model and that with no phenotype differences (Model B) provides evidence that there are phenotype differences between correlations (see Methods). The 95% highest-density posterior (HDP) regions of each of the phenotype-level parameters are then inspected to determine in which parameters the phenotypes differ. The 95% highest-density posterior regions are disjoint for the following parameters: *μ_α_, μ_γ_,μ_A_* and *σA* (Figure 7). This can be interpreted biologically as follows: there is evidence that wild type and *β*2( *TG)* spontaneous retinal activity differ in the following respects: the baseline level of correlated firing (lower in *β2(TG))*, the extent of the waves - related to the rate of decay of correlations *(β2 (TG)* waves are truncated), the variance in the amount of correlated firing outside of waves (lower in *β2 (TG))* and the inter-recording variance in the amount of correlated firing outside of waves (lower in *β2(TG)).*

## Modelling data from Blankenship et al. [5]

To test the generality of our hierarchical Bayesian model, as well as analysing the data from Xu et al. [69], we then tested how well the model could be applied to another dataset without further modification of the model. As a validation set, we therefore chose the dataset by Blankenship et al. [5] primarily because it has three phenotypes (one wild type and two mutant genotypes) rather than just two.

### Modelling process

The modelling process and the data set have been described in Methods. The best-fitting data-distribution was found to be the exponentially modified Gaussian (EMG) distribution with the Gumbel distribution being the next best fit (Figure 9). The Gumbel distribution was chosen as the data-generating distribution on the basis of the practical considerations described in Methods: the Gumbel distribution has two parameters and the EMG has three. The third parameter of the EMG shows great variation and any distance-dependence function which was fitted had large errors and a large range of distance-dependence parameters which made sampling difficult.

The Gumbel distribution has a location parameter *u* and a scale parameter *v* and the distance-dependence functions. As with the data from Xu et al. [69], F3 was the best fit to the distance-dependence of v. F9 was the best fit for u, with F1 being the next best fit (Figure 9). F1 was chosen as the distance-dependence function since it is preferred as all its parameters have a intuitive biological interpretation and the range of fitted distance-dependence parameters are large for F9 (which resulted in problems finding suitable initial values) and small for F1.

**Figure 9:**
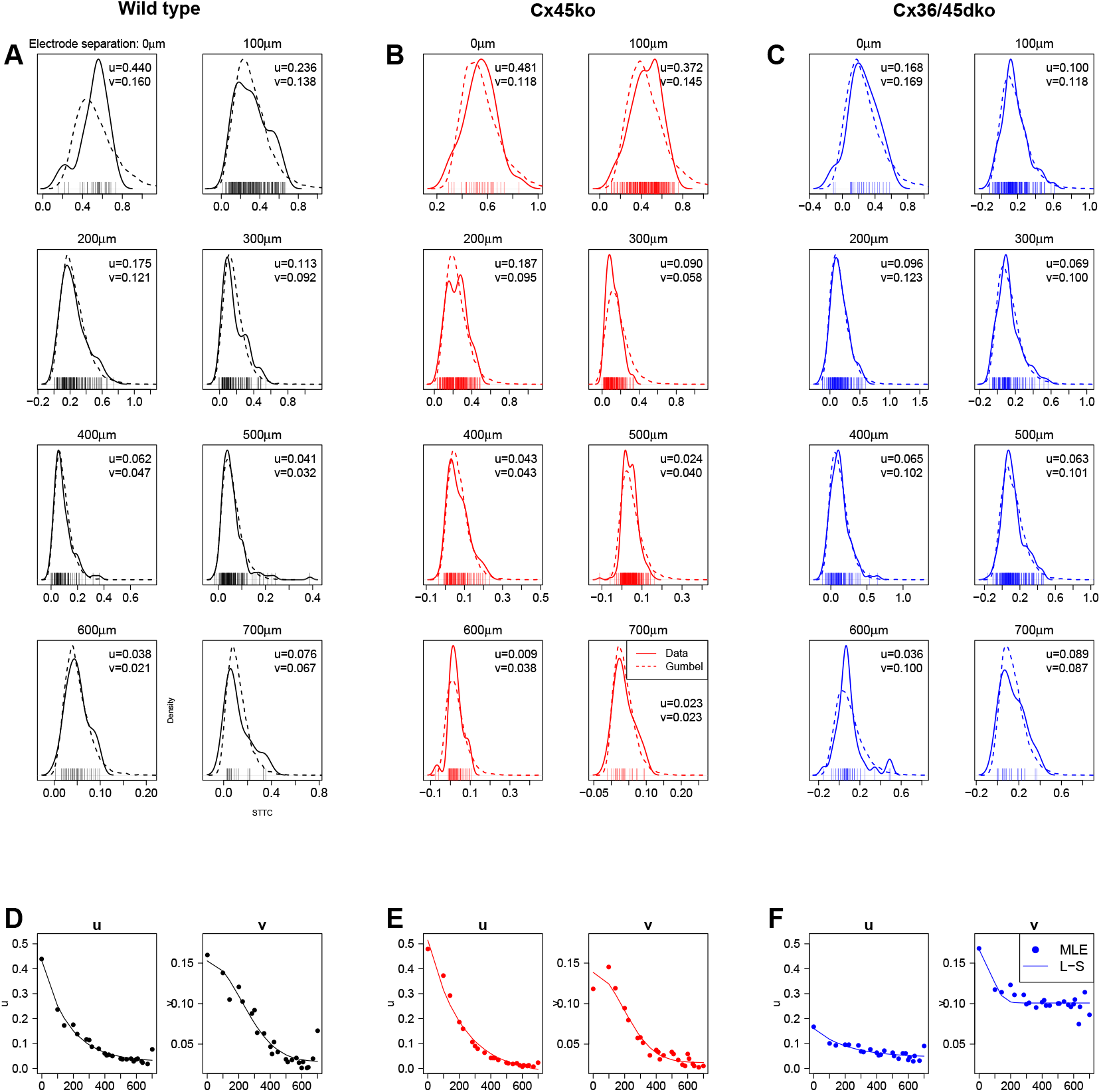
Maximum likelihood (ML) fits of the data-generating distribution and least-squares (LS) fits of the distance-dependence functions to data from Blankenship et al. [5]. Panels as per Figure 6 but with three pheonotypes: one wild type (A,D), one Cx45 knockout (B,E) and one Cx36/45 double knockout (C,F).

The full model is therefore as it was for the data from Xu et al. [69] with the only difference being that the phenotype-level parameters exist for each of the three (as opposed to two) phenotypes considered. The bounds on the phenotype-level parameters are the same as those used for that model.

### Model Assessment

The results of the modelling process are the posterior distributions of all the model’s parameters. Since we are interested in phenotype-level differences we only present the posterior distributions of the phenotype-level parameters (Figure 10).

**Figure 10:**
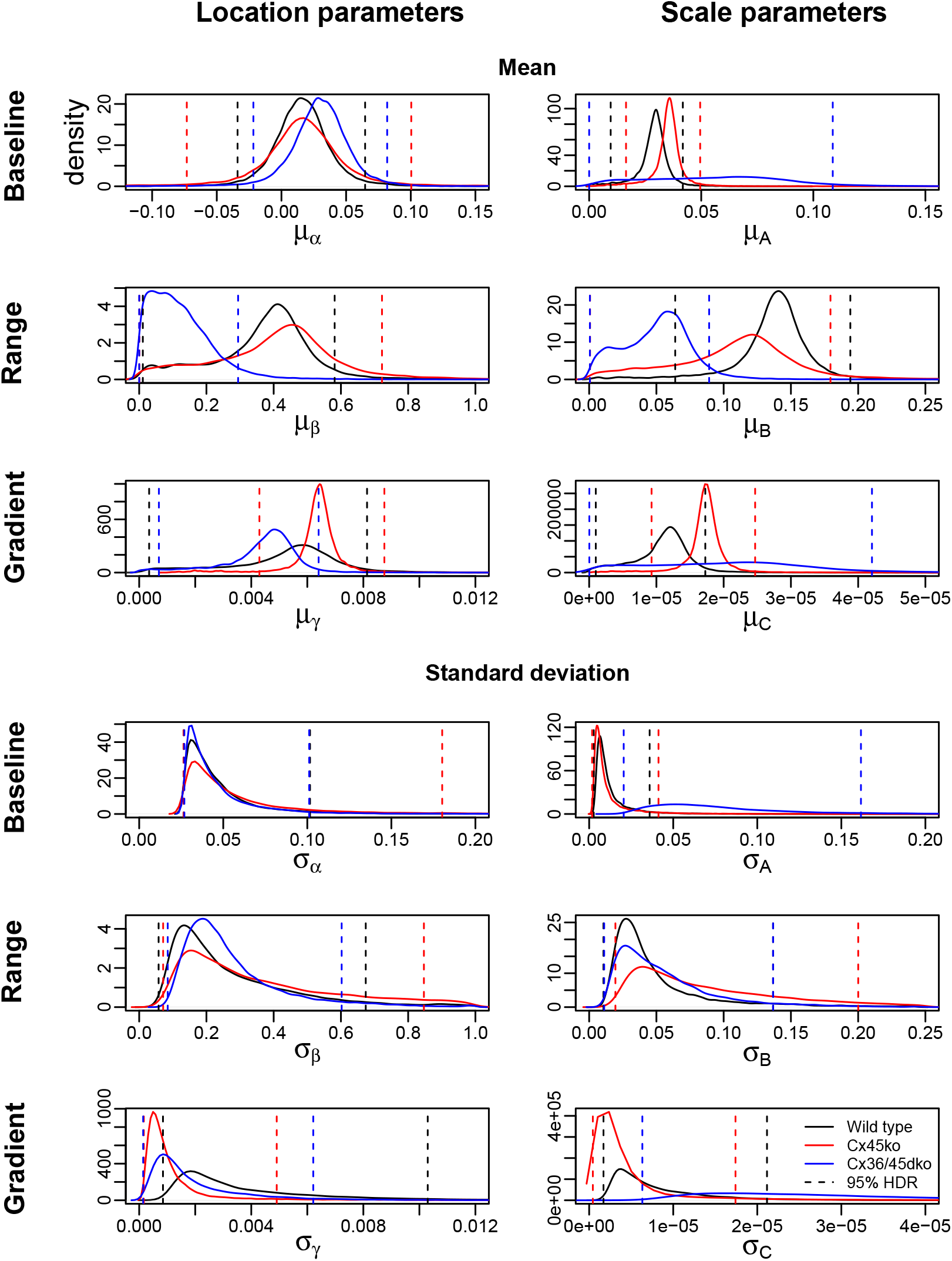
Posterior distributions of the phenotype-level parameters for Model F of data from Blankenship et al. [5]. Panels are as in Figure 7.

Our model (Figure 11A and B) fits the data well; as with the data from Xu et al. [69] the main cause of discrepancy is the fact that sampling from the model occasionally produces STTC values which are ≥ 1.

**Figure 11:**
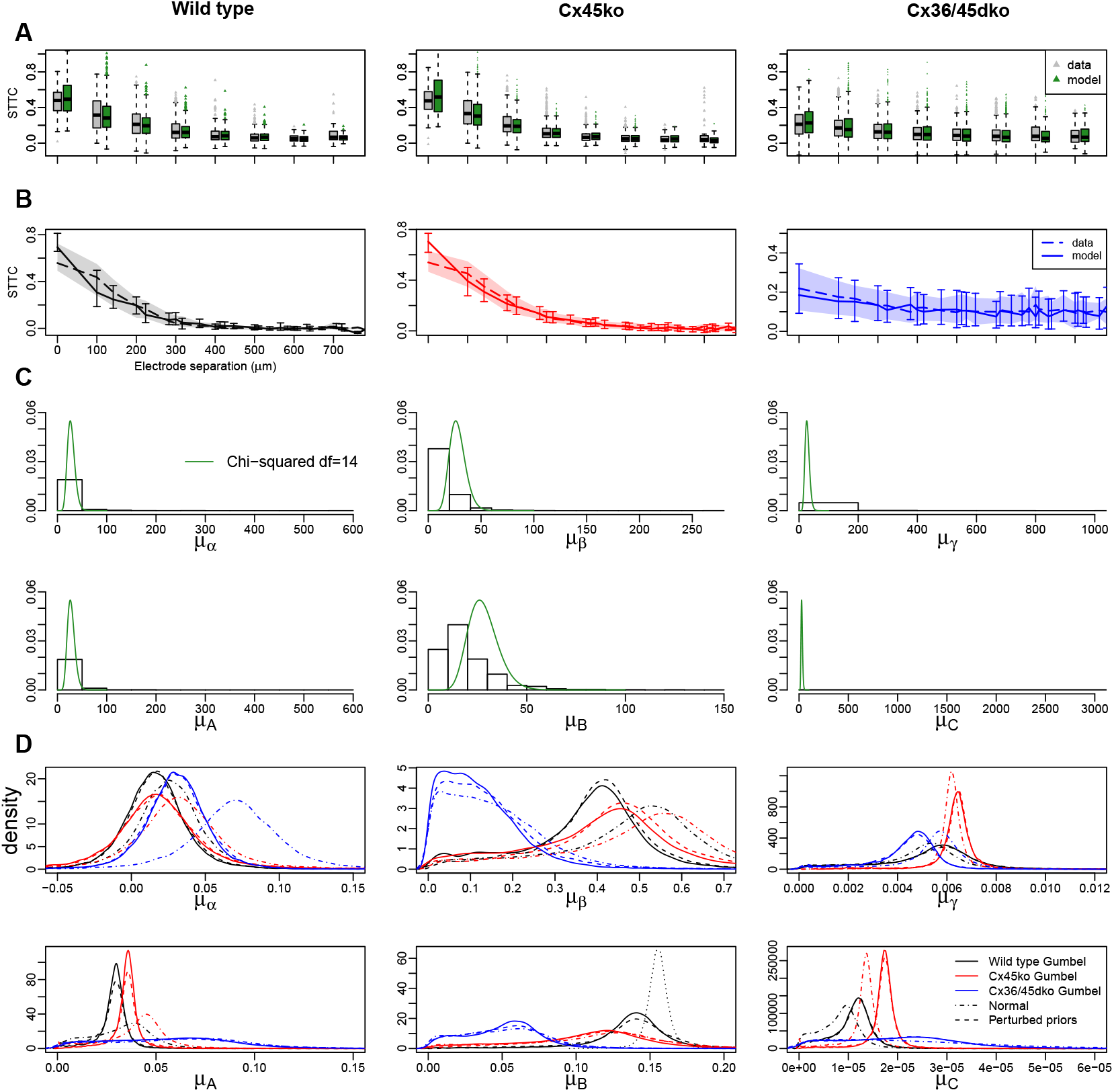
Assessment of Model F of data from Blankenship et al. [5]. Panels are as described in Figure 8, with the following differences. A: Data is shown for each of the three phenotypes (left - wild type, center - Cx45ko, right - Cx36/45dko). B: Data is shown for three recordings, one from each phenotype (order as in A). C: The theoretical distribution of the PDM is χ^2^ with 14 degrees of freedom (as two recordings were removed as outliers). D: Prior distributions for the model with perturbed priors were assumed to be normally distributed [*μ_α_ ∼ N(0.05, 0.5), μ_β_ ∼ N(0.75, 0.5), μ_γ_ ∼ N(0.1, 0.25), μ_A_ ∼ N(0.3, 0.25), μ_B_ ∼ N(0.4, 0.25), μ_C_ ∼ N(0.1, 0.1)]*.

Assessment of the model’s performance at the recording level (Figure 11C) shows that the distribution sampled from the data is over-dispersed compared to the theoretical distribution. The posterior distributions of the phenotype-level parameters are much less localised for this data set than those of the data from Xu et al. [69] (compare Figures 7 and 10) which means that "extreme" values are more likely to be drawn which can give large values of the PDM. While this is not ideal, it is not overly concerning since the model’s ability to replicate the data is good.

Assessment of the model’s robustness to both its prior and its assumptions (Figure 11D) show that the conclusions do not change, even with strong perturbations.

**Table 6.** WAIC values for each model, listed in increasing order, of the Blankenship et al. data [5].

| Model | WAIC |
| --- | --- |
| B | -95166.33 |
| F | -95166.08 |
| G | -95164.94 |
| A | -79468.68 |
| C | -79279.73 |

Assessment of the importance of including phenotype-level differences and recording-level differences is performed using the WAIC (Table 6). Model B has the lowest WAIC, implying it is the most parsimonious fit, but the difference between its WAIC and that of Model F is too small (0.25) to be considered indicative of a preferred model. The same is true of the difference in WAIC between models F and G (1.14). This demonstrates that the differences between recording are key to explaining STTC variance in this data set, rather than the differences between phenotypes. This is further backed-up by the fact that there is a large improvement in fit between Models A and C (which have no recording-level differences) and Models B, F and G (which do have recording level differences). In addition the posterior distributions of Models A and C are highly localised compared to those of Models B, F and G (Table 7) meaning that these models would be over-confident in the location of the parameter.

**Table 7.**
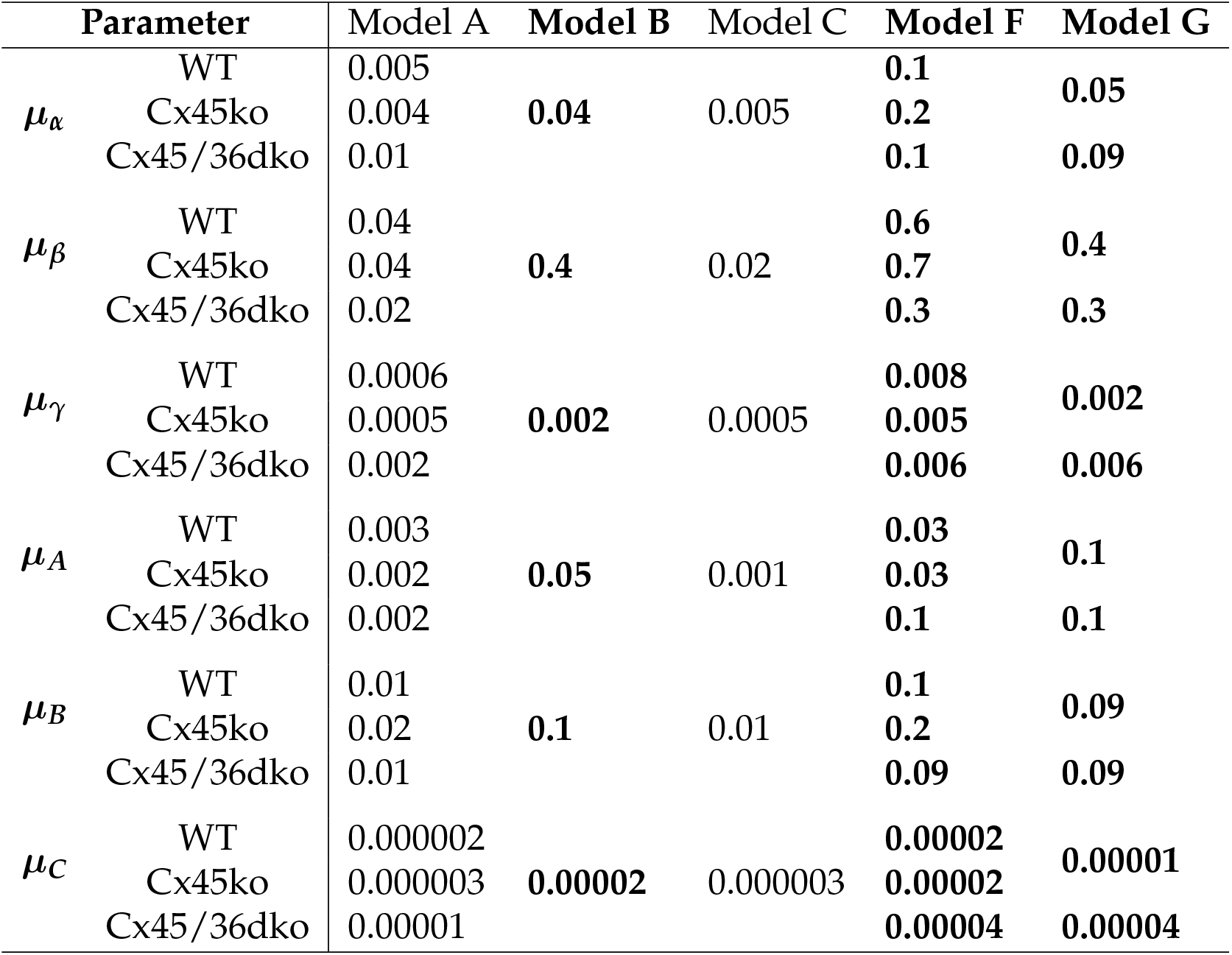
Widths of 95% highest posterior density (HPD) regions for Models A-G for the Blankenship et al. data. Data are shown as per Table 5.For models (A and F) where the parameter is phenotype-level dependent, the HPD widths for both wild type, Cx45ko and Cx45/36dko are given. Model G assumes that wild type and Cx45ko are indistinguishable, so only two phenotypes appear. Models (B, F and G) where there is recording-dependence appear in bold.

### Assessing evidence for/against differences between phenotypes

The difference in WAIC values between Models B, F and G are so small as to provide no evidence that there are differences between phenotypes. This is further supported by the fact that the 95% highest-density posterior (HDP) regions are not disjoint between phenotypes for any of the phenotype-level parameters so we conclude that there is no evidence for phenotype-based differences in this data set. This goes counter to the previous conclusions, based upon visual inspection only, that the wild type and Cx45ko correlations are similar, and that the Cx36/45dko is distinct from the two other distance-dependent correlations.

## Discussion

This work has investigated the methods of inference used to decide if the correlation-distance relationship of spontaneous retinal activity differs between experimental conditions. Less than half of all publications which contained a correlation-distance plot tested for the significance of their conclusions and the remaining publications used standard frequentist tests. We argued that these approaches are problematic and proposed a framework for Bayesian modelling and inference on the correlation-distance relationship. To demonstrate its use we applied it to two data sets Xu et al. [69] and Blankenship et al. [5]. We find evidence that the distance-dependence of correlations differs between the wild type and *β*2(TG) phenotypes in the data from Xu et al. [69] (this is in line with previous conclusions based on visual inspection). We find no evidence that the distance-dependence of correlations varies between the three phenotypes (wild type, Cx45ko, Cx36/45dko) considered in the data from Blankenship et al. [5]. This runs counter to previous conclusions based on visual inspection (that there were differences in correlations between phenotypes) and demonstrates the need for thorough statistical analysis of the distance-dependence of these correlations in order to draw robust conclusions.

### Insights from analysis of data

Our analysis provides evidence that wild type and *β*2(TG) phenotypes from Xu et al. [69] differ in the extent of the wave (rate of decay of correlation), the level of correlated firing outside of waves and the variation in correlations outside of waves. The data from Blankenship et al. [5] showed no evidence for differences between the three phenotypes (wild type, Cx45ko and Cx36/45dko): although the posterior distributions of Cx36/45dko were offset from the other two phenotypes (in general), the long tails prevented differences being found by our ad-hoc test which compared overlap of highest posterior density regions.

The results from Xu et al. [69] data broadly confirm the intuition from the correlation-distance plots but the results from Blankenship et al. [5] demonstrate that inspection of the correlation-distance plots can be misleading: the summary statistics (median and IQR) mask the large inter-recording variance and there is a tendency to concentrate on the median values and ignore the effects of variance which leads to the conclusion that there are differences between Cx36/45dko and the other two phenotypes. Our analysis framework models the full complexity of the data and demonstrates that these differences are not significant.

The long tails of the posterior distributions in the Blankenship et al. [5] data are an impediment to our ad-hoc test since HPD regions overlap despite clear differences in the posteriors. These tails may be due to a greater amount of variance across the data in all respects (making it hard to localise posteriors) or due to the small number of recordings per phenotype Xu et al. [69] has 13 and 17 recordings per phenotype, Blankenship et al. [5] has 4,5 and 6).

The Cx36/45dko mutant shows defects in eye-specific segregation, but the analysis did not show significant differences in correlations between this and wild type. Three possible explanations are: firstly, that features which are not measured in this model are responsible for the formation of eye-specific segre gation. Secondly, that the Cx36/45dko mutant is different from the other two, but the amount of data available is insufficient to give sufficiently localised posteriors to distinguish this. Thirdly, that the weight of evidence which we require to demonstrate that there is a difference is more stringent than that which is required for eye-specific segregation (i.e. the biological system is more sensitive than our framework). Without more data it is difficult to be more specific.

### The choice of model

The framework is flexible and can be used to fit models to other data sets. The same model was fitted to both data sets considered here which implies that it may capture some inherent features of the correlation-distance relationship and is a reasonable starting point for performing inferences on other data sets (although the assumptions should be checked and the model altered if necessary).

It is not surprising that exponential decay was chosen as the distance-dependence function: it is a relationship which occurs frequently in the biological world. The choice of the Gumbel distribution is unlikely to have any physical relevance to the system as, despite it fitting the data well (and being the most pragmatic choice), there are discrepancies between it and the data (e.g. it has a fixed skew, but the skew of the data varies).

The form of the fitted model was useful as it has bounded parameters making computational time reasonable and convergence unproblematic. This may not be the same for other models. Using the STTC also helped bound parameters, but the approach can be used with any measure of correlation. The model and model fitting is flexible: the number of variables can be increased to consider any number of phenotypes, ages and experimental conditions.

This work highlights the importance of using hierarchical modelling to capture inter-recording variations within phenotypes. Models which ignored these (A and C) were poor fits to the data with over-confidence in the parameter locations and the model with recording-level and no phenotype-level differences (B) fit the data almost as well as (data from Xu et al. [69]) or as well as (data from Blankenship et al. [5]) the full model. The distance-dependence of the correlations is key to their differences and so the results from our framework are more credible than those which ignore this and inter-recording variation or inspection alone. The framework has two further advantages: Bayesian analysis is more informative than a hypothesis test alone (posterior distributions as opposed to point-estimates) and the fact that the models parameters have a physical interpretation so the features of the correlation which differ can be investigated, as opposed to just if they differ in some (unknown) respect. We believe that the method is sufficiently intuitive and pragmatic to be useful in practice and that the improvement in the results is worth the extra complexity.

It should be noted that this method assesses the evidence for mathematical/statistical differences which is different from assessing evidence for biological significance: the systems sensitivity to changes in correlation is unknown and our definition of significant could be more or less stringent than that which the visual system can discern. This is true for much of experimental biology [32, 41] evidence for a difference between phenotypes does not imply that this difference has a biological significance but it demonstrates that it exists and is unlikely to be caused by chance.

### Wider applicability

In addition to a Bayesian framework, the methods of inference used in this paper differ from those previously used in that we model the variation between recordings of animals of the same phenotype as opposed to pooling across phenotypes. Data where multiple measurements are taken from the same object are described as "nested". Nested data is very common in neuroscience (e.g. neuron morphology studies) and also in the wider biological sciences (e.g. in genetics, families are not genetically independent and in medicine, patients can be considered as nested by hospital). In a recent large literature review of eighteen months of all molecular, cellular and developmental articles in Science, Nature, Cell, Nature Neuroscience and (every first monthly issue of) Neuron, at least 53 % of 314 articles used nested data, but all of these studies used conventional statistical analyses (e.g. t-test on pooled data) which failed to take account of the nested nature of the data [1].

Pooling nested data violates the common statistical assumption that obser vations are independent and leads to a large increase in the false positive rate of standard statistical tests [1]. This is a contributing factor to the recently-raised high-profile concerns about the contamination of neuroscience literature by false positives [7,29,44]. Frequentist hierarchical (also called "fixed-effects") models can be used to accommodate nested data and analogies to the standard frequentist tests can be used which may be adequate for some data sets, however we believe a Bayesian approach to be more appropriate for our data set due to its flexibility and the very natural way of incorporating multiple-levels.

The expense and time involved in neuroscience investigations means that it is advantageous to make as many recordings as possible from each subject and the number of techniques where this is common practice is large. This is decidedly not limited to analysis of MEA data, nor electrophysiological recordings. Aarts et al. [1] identify the following non-exhausting list of techniques where nested data is frequently collected: analysis of immunofluorescence signal intensity in slices, optogenetics, super-resolution microscopy, immunogold cytochemistry and optophramcology. There is, in general, a caveat that the number of recordings from each unit must be sufficiently large for multi-level recording to be robust (otherwise these techniques are not helpful). As a rule of thumb this is given to be about five observations per unit [37, 51], which is much smaller than the number of observations per unit in the data sets considered here (of the order 1,000).

## Conclusions

We believe that the framework outlined in this work offers a substantial improvement on previously used methods of inference and is more robust and credible than frequentist hypothesis tests as its specifies a model for the correlation-distance relationship which was previously ignored and as it accounts for the (often large) inter-recording variations within phenotypes. As it is able to highlight features of the correlation-distance relationship which differ between phenotypes it should help form a more detailed understanding of the role of correlations in map development.

## Acknowledgements

The authors thank the Wellcome Trust (SJE; grant number 083205) and EPSRC (CSC) for financial support.

## Availability of research data

The models here have been written in the R programming language, and make extensive use of Stan [56]. The retinal wave recordings that were analysed here came from [21], and are available at http://dx.doi.org/10.5524/100089.

